# Draft genome sequence of the glasshouse-potato aphid *Aulacorthum solani*

**DOI:** 10.1101/2024.11.12.623246

**Authors:** Joseph Torres, Paula Rozo-Lopez, William Brewer, Omid Saleh Ziabari, Benjamin J. Parker

## Abstract

*Aulacorthum solani* is a worldwide agricultural pest aphid capable of feeding on a wide range of host plants. This insect causes mechanical damage to crops and is a vector of plant viruses. We found that the publicly available genome for *A. solani* is contaminated with another aphid species, and we produced a new genome using a barcoded isogenic laboratory line. We produced Oxford Nanopore and Illumina reads to generate a draft assembly, and we sequenced RNA to aid in the annotation of our assembly. Our *A. solani* draft genome is 671 Mbp containing 7,020 contigs with an N50 length of 196 kb with a BUSCO completeness of 98.6%. Out of the 24,981 genes predicted by E-GAPx, 22,804 were annotated with putative functions based on homology to other aphid species. This genome will provide a useful resource for the community of researchers studying aphids from agricultural and genomic perspectives.

## Introduction

The glasshouse potato aphid, *Aulacorthum solani* (NCBI Taxonomy ID: 202456; also referred to as the foxglove aphid), is a polyphagous species with one of the broadest host plant ranges of any aphid. This species is an agricultural pest of crops such as tomatoes, peppers, tobacco, cucurbits, and legumes (1). It is a particularly important pest of greenhouse crops in North America and the UK (2). In addition to direct feeding damage that causes discoloration and deformation of leaves, *A. solani* aphids can also vector several plant viruses (3).

More broadly, aphids are important study organisms for addressing questions in evolutionary biology, developmental biology, and microbial symbiosis (4, 5). However, only a small percentage of the 5,000 estimated aphid species have available whole genome sequences in NCBI and/or AphidBase (6). *A. solani* sits within the tribe Macrosiphini along with key species including the model pea aphid (*Acyrthosiphon pisum*) (7), making it a critical species for comparative studies.

We uncovered issues with the previously published reference genome for *A. solani* (NCBI accession ASM852887v1); (8). We show that this assembly is likely contaminated with sequences from multiple aphid species, and that some or all of the genome comes from another aphid species. This misidentification is leading to issues in the literature. For example, a recent attempt to resolve the aphid phylogeny based on ultra-conserved elements used the misidentified ASM852887v1 genome. This led to an incorrect placement of *A. solani* in the aphid phylogeny (9), complicating taxonomic and evolutionary studies of aphids.

Here we describe a high-quality genome assembly for *A. solani* using an isogenic laboratory line. We generated a draft genome using Oxford Nanopore and Illumina reads, and sequenced RNA to aid in genome annotation. Our draft genome is 671 Mbp containing 7,020 contigs with an N50 length of 196 kb, and BUSCO scores and other analyses suggest the genome is highly complete. Our annotation contains 24,981 genes, and we use this to compare *A. solani* to the closely related *A. pisum*.

## Methods

### Aphid collection and colonization

We collected asexual winged and wingless female *Aulacorthum solani* adults from garden tomato plants in Knoxville, TN, USA, in May 2021. To establish a colony of *A. solani* in the laboratory, we used a single asexual female. After colonization, we maintained this line on fava bean plants at 20ºC 16L:8D. To validate our taxonomic identification, we used COI barcoding (LCO1490 5’-GGTCAACAAATCATAAAGATATTGG-3’ and HCO2198 5’-TAAACTTCAGGGTGACCAAAAAATCA-3’), sanger sequencing, and comparisons of our COI sequence to the Barcode of Life Data System (10). The BOLD System database allows animal identification (https://v3.boldsystems.org/index.php/IDS_OpenIdEngine) and bases species matches using Neighbor-Joining trees constructed from the top hits. Our partial COI barcode sequence was uploaded to NCBI with accession number PQ361290.

### Species ID for the existing *A. solani* assembly

We analyzed the COI sequence found in the putative glasshouse-potato aphid assembly (accession ASM852887v1) to troubleshoot problems with this genome. We downloaded *A. pisum* COI accessions from the Barcoding Life Database (record ID “ACEA116-14”) and used the sequences to BLAST the genome assembly. We then took the top alignment that spanned the entire region and queried it in the Barcoding Life Database as above.

### DNA extraction and sequencing

We pooled seven genetically identical adult unwinged aphids cultivated in the laboratory and isolated genomic DNA (gDNA) using Bender-buffer and an ethanol precipitation (11). We then sheared the gDNA to approximately 20kb fragments using Covaris G-tubes (Covaris LLC., Woburn, MA, USA) at 4200 RMP for 1 minute, followed by tube inversion. Part of the sample was set aside for Illumina sequencing. For Oxford Nanopore library preparation, we used the NEB Next PPFE repair kit with Ultra II end prep reaction (New England Biolabs, Ipswich, MA, USA) under recommended conditions and Nanopore ligation sequencing kit SQK-LSK110. For sequencing, we used a Nanopore R9.4.1 (FLO-MIN106D) flow cell and a MinION MIN-101B sequencing device (Oxford Nanopore Technologies, Oxford, UK). We ran the flow cell for 24 hours, followed by a wash with Flow Cell Wash Kit (EXP-WSH004); we then reloaded the flow cell with a second library prep and ran the sequencer for an additional 48 hours. We stopped the second sequencing run at 24 hours (∼11 Gbps of sequencing). Raw nanopore sequencing reads are available in the NCBI Sequence Read Archive under Bioproject ID PRJNA1156622 with Biosample accession SAMN43496284 and Run ID SRR30661341. In addition, we conducted DNA Illumina sequencing at Novogene (Novogene Corporation Inc., Sacramento, CA, USA) using DNA extracted as above. We generated 15.5 billion base pairs with 150 bp paired-end reads on an Illumina NovaSeq platform. Illumina reads are available in the NCBI Sequence Read Archive with Biosample accession SAMN43799130 and Run ID SRR30686038.

### RNA extraction and sequencing

We homogenized pools of five adult aphids with a pestle in 800 µL of TRIzol (Invitrogen; Thermo Fisher Scientific, Inc., Waltham, MA, USA) and extracted total RNA using Chloroform and isopropanol precipitation. We used the Zymo RNA Clean & Concentrator kit (Zymo Genetics Inc., Seattle, WA, USA) to improve the purity and to remove DNA using DNAse I. We then performed transcriptome Sequencing at Novogene (Novogene Corporation Inc., Sacramento, CA, USA). Library preparation was conducted using poly-A selection. The libraries were sequenced to approximately 3.4 billion base pairs (bp) per sample with 150 bp paired-end reads on an Illumina NovaSeq platform. Raw reads were deposited into the NCBI Sequence Read Archive under BioProject ID SRP532563 with BioSample accession SAMN43784434 and Run ID SRR30675520.

### *A. solani* whole genome assembly

We used Guppy (Oxford Nanopore Technologies) for base-calling and quality trimming raw reads (12). We assembled Oxford nanopore reads using Canu v. 2.0 with recommended parameters and a predicted genome length of 300Mb (13). We then mapped Illumina reads to the assembled contigs using BWA (14). We used these alignments to polish assembled contigs with Pilon for two rounds (15) and then assessed with BUSCO v. 5.7.0 using Hemiptera as the Busco lineage parameter (16). We removed haplotigs with PurgeHaplotigs by mapping the contigs against the original Oxford nanopore reads and selecting the appropriate valley points as shown in the coverage histogram (Figure S1) (17).

### Contig purging

We removed all contigs that matched a phylum other than Arthropoda, including from *Proteobacteria*, using the Blobtools visualizer after BLAST searching the contigs against the NCBI nucleotide NR database with the BLASTplus command line interface (Figure S2) (18, 19). In addition, we manually removed 2,058 contigs based on mapped Illumina read depth and an average short read coverage below 14x, as the standard deviation was higher than the average coverage. The remaining contigs were assessed again with BUSCO. The *A. solani* genome is available in NCBI with BioProject ID PRJNA1156622 accession JBHJKM000000000.

### *A. solani* genome annotation

We used RNAseq data to annotate the assembled contigs using NCBI’s alpha-phase E-GAPx v. 0.2 annotation pipeline. We used the resulting gene predictions for automated functional annotation with the EnTAP pipeline using the default Eggnogmapper with homology searches against the NCBI protein NR database, the Uniprot Swissprot database, the Uniprot Tremble database, and the ref-seq invertebrate database with bacteria set as the contaminant flag (20). We then submitted the predicted genes to BLAST-koala to obtain up-to-date KEGG ortholog terms (21). We compared the pathway modules to the pea aphid *A. pisum* AL4f, Project ID PRJNA547584, Biosample SAMN10253041. The individual commands and slurm scripts as well as the KEGG assignments are available on GitHub (https://github.com/Artifice120/Aulacorthum_solani_genome_assembly_pipeline/tree/main).

## Results

### The existing *A. solani* assembly is likely *Brachycaudus helichrysi*

BLAST results of the pea aphid COI sequence from the BOLD database against the ASM852887v1 genome assembly hit the PVMI01058906.1 scaffold from base pairs 9069-9726 with a 92.9% identity. We found that this region of the genome matched *Brachycaudus helichrysi* as a top hit with a 100% match to the BIN barcode in BOLD systems. For our genome, we only used aphids from an isogenic colony (AS13) that has been maintained in the laboratory for multiple generations and confirmed the species identification by COI Barcoding. Furthermore, after genome assembly, we reconfirmed the presence of the COX1 barcode identification for *A. solani* in contig tig00020064 (9727-10437) with a 100% match to the *A. solani* BIN barcode in BOLD systems.

### *A. solani* whole genome assembly

We obtained a total of 4,873,637 nanopore reads (at an average length of 6.7kb) and 10,344,638 Illumina reads (PE 150bp). After assembly, haplotig purging, polishing, removal of plant and bacterial contigs, and contig bubble resolution, our assembly contained 7,020 contigs with an N50 length of 196 kb and a total length of 671 Mbp. The assembly has a BUSCO completeness score of 98.6% (n= 2,510) and a duplication score of 14.4% (Figure 1). The genome size is slightly larger than closely related species including *A. pisum* (514.2 Mbp) but within the range of aphid genomes (317.1 Mbp – 711.3 Mbp estimated by flow cytometry (22)). GC content at 30.0% was similar to other aphid species (*A. pisum:* 29.6% (23); *M. persicae:* 30.1% (24)).

**Figure 1.**
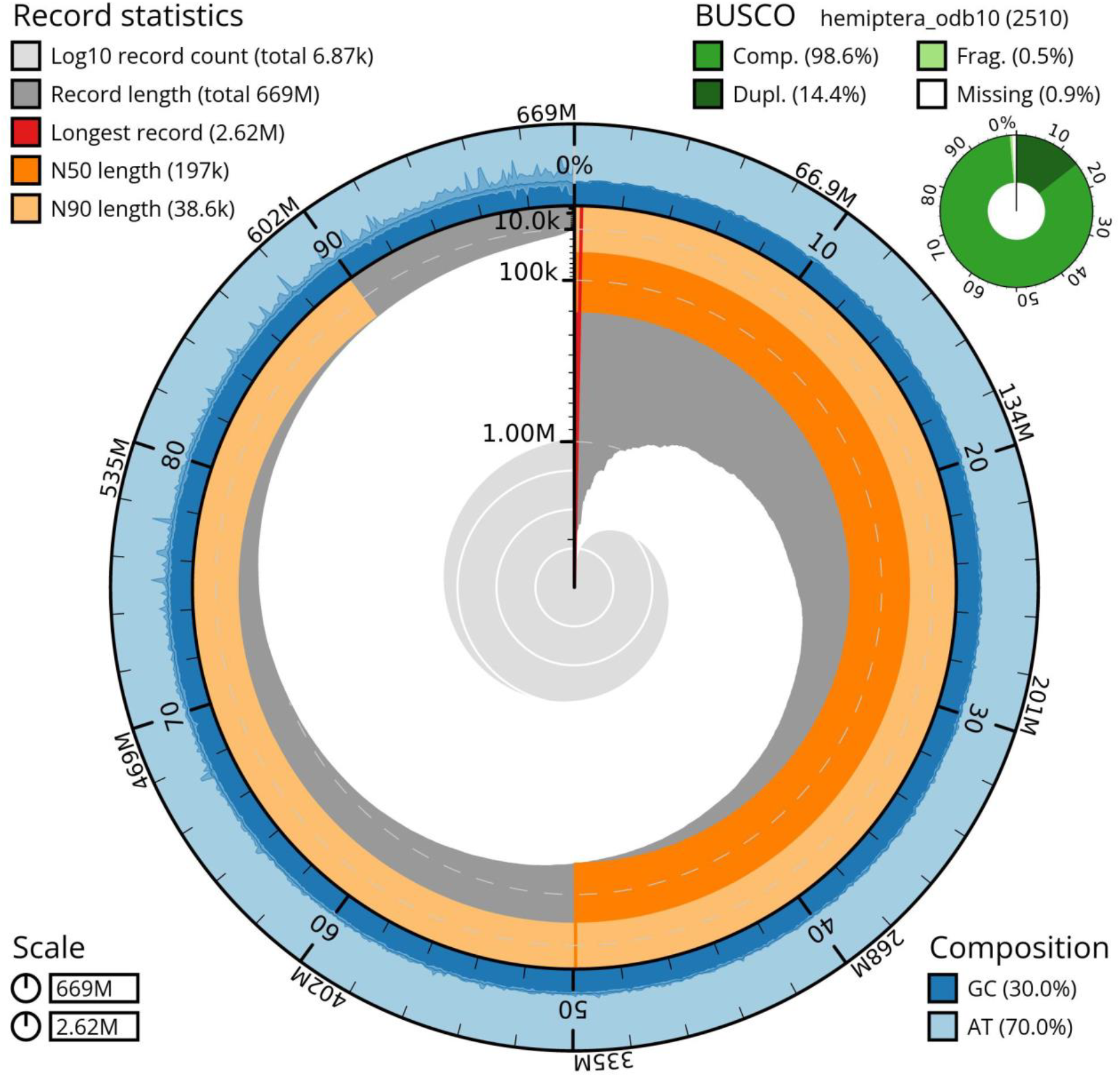
Snail plot summary of assembly statistics for *A. solani* assembly. The main plot is divided into 1,000 size-ordered bins around the circumference of the snail plot with each bin representing 0.1% of the 670,586,136 bp assembly. The distribution of the sequence lengths is also shown in dark grey with the plot radius scaled to the longest sequence in the assembly (2,615,422 bp, shown in red). The orange and pale-orange arcs represent the N50 (195,973 bp) and N90 (38,241 bp) sequence lengths. The pale grey spiral shows the cumulative sequence count on a log scale with the white dotted lines showing successive orders of magnitude. The outermost pale-blue area on the outer edge of the plot represents the distribution of the GC and AT percentages. A summary of the complete, fragmented, and duplicated BUSCO genes in the hemiptera_od10 set is shown in the top right corner.

In addition to contigs with homology to the aphid obligate symbiont *Buchnera aphidicola*, we removed contigs with homology to the alphaproteobacteria *Wolbachia*, suggesting that our aphid line is infected with *Wolbachia*. Analysis of these contigs yielded a BUSCO score of 37.3% complete using the Rickettsiales_odb10 database, suggesting the microbes were not at high enough density in our aphid sample to produce a useful genome assembly for this bacterium under these conditions.

### *A. solani* genome annotation

24,981 genes were predicted by E-GAPx based on protein alignments and HMM models appropriate to the taxonomy as well as RNA-sequencing data. We found 93.4% of functional annotation assignments are within the Arthropoda lineage, with the highest homology towards *A. pisum* (pea aphid) (Figure S3). This was visualized further with a synteny plot showing the mapped regions between the pea aphid reference genome and this *A. solani* draft genome (Figure S4). Many of the annotated genes had putative functions such as detoxification genes, salivary genes, virus transmission genes, transcription factor genes, and mitochondrial genes (Table 1), which are also found in other aphid species such as *A. pisum, Metopolophium dirhodum, Myzus persicae, Aphis craccivora*, and *Aphis glycines*. Out of the 24,981 genes predicted by E-GAPx, 15,332 were annotated with families were different from A. pisum with 22,780 annotated genes total from the aphid superfamily. In addition, we found 98 total cytochrome P450 oxidase genes that were distributed across 30 families (Figure S5).

**Table 1.** Number of predicted *A. solani* genes with putative functions based on the reference protein databases (NCBI BlastX ‘Nr’ (non-redundant), UniProt Swissprot, UniProt tremble, and the ref-seq invertebrate database).

|  |  |
| --- | --- |
| i. Defensive and detoxification genes |  |
| a) Cytochrome P450 | 98 |
| b) Glutathione S-transferase | 29 |
| c) ABC transporter | 66 |
| d) Carboxyl/cholinesterase | 41 |
| e) UDP-glucosyltransferase/UDP-glucuronosyltransferase | 123 |
| f) Cuticle protein | 91 |
| ii. Salivary genes and chemoreceptors |  |
| a) Glucose dehydrogenase | 36 |
| b) Sugar/glucose/inositol transporter | 41 |
| c) Apolipophorin | 2 |
| d) Salivary gland/protein/peptide | 8 |
| e) Secretory protein | 10 |
| a) Gustatory receptor | 215 |
| b) Odorant receptor/protein | 124 |
| iii. Insecticide resistance genes |  |
| a) Acetylcholine receptor/esterase/transporter | 21 |
| b) Sodium exchanger/symport/channel/neurotransmitter/transporter | 100 |
| c) GABA transporter | 2 |
| d) Multi-drug resistance protein | 30 |
| iv. Virus transmission genes |  |
| a) Dynamin | 37 |
| b) Serine protease inhibitor | 12 |
| c) Vesicle transport/trafficking | 10 |
| d) Endocytosis | 8 |
| e) Exocytosis | 4 |
| v. Transcription factors | 379 |
| vi. Mitochondrial genes | 599 |

### Differences with the pea aphid genome

We identified several potentially significant differences between the *A. solani* and *A. pisum* genomes. First, *A. solani* appears to have an expanded number of odorant receptors, with 17 different odorant receptor families present in our annotation compared with 9 in the pea aphid genome. Next, KEGG annotations highlighted three complete pathway modules that are present in *A. solani* that are missing from pea aphids (M00131, M00132, and M00415) (21). Specifically, M00131 and M00132 are part of Inositol phosphate metabolism, with the first incorporating myo-inositol and the second incorporating phytate into Inositol phosphate metabolism. The pathway module M00415 corresponds to fatty acid elongation in the endoplasmic reticulum while the pea aphid KEGG annotations only show complete modules with fatty acid elongation in the mitochondria.

## Discussion

We generated an accurate draft genome assembly for *A. solani*. Our analysis of single-copy orthologs and complete BUSCOs (98.6%), suggests that our *A. solani* draft genome assembly is highly complete. We found evidence of infection of this aphid with *Wolbachia* though we could not assemble a genome for this symbiont from our data, but this result does add to the growing number of aphid species that have been identified as hosting *Wolbachia* suggesting it is more common among aphids than was previously acknowledged. Future work to reduce duplication in the genome and chromosome-scale genome scaffolding would improve this genome assembly, and additional coverage could produce sequences for *Wolbachia* and any other microbial associates of this aphid.

A key difference between *A. solani* and the model pea aphid appears to be the incorporation of myo-inositol and phytate into inositol phosphate metabolism. Pea aphids have been shown to not absorb myo-inositol or its free cyclitol form (phytic acid); the concentration of cyclitols in pea aphid honeydew is higher than the concentration in plant stems and leaves (25). An intriguing possibility is that these metabolic differences contribute to the broad host range of *A. solani*. Host plant range differences could also be linked to the expanded repertoire of odorant receptors found in *A. solani*, and future work could focus on the genomic basis of herbivore generalism given the broad host range of this insect.

Our work also corrects the record with respect to the existing ASM852887v1 assembly for *A. solani*. BLAST results suggest this assembly was either misidentified or was contaminated by multiple aphid species, including *Brachycaudus helichrysi*. We ensured our barcode-confirmed *A. solani* isogenic line had no contamination from other species at every step of the genome assembly process. Identification of aphids is difficult as many species exhibit significant similarities in their morphology. Misidentification is often compounded by phenotypic variation within a single species, making accurate identification challenging without specialized mounting techniques or molecular methods like DNA barcoding. In addition, multiple species can be found on the same host plant species, and there is a need to carefully colonize isogenic lines from single parthenogenic aphids in the laboratory before generating sequencing data originated from pooling individuals. The genus *Aulacorthum* is especially challenging since there are likely cryptic species groups, species placements shared with *Acyrthosiphon*, and poor coverage of barcode reference sequences for non-agriculturally relevant species.

Along with quality assemblies from a growing number of aphid species, the *A. solani* draft genome provides a valuable tool for studying insect diversification and evolution, plant-insect interactions, and aphid-virus interactions. Future work using this genome will increase the power of aphid systems for addressing basic biological questions, and could have practical application in the control and management of this pest species.

## DataAvailability

This project can be found under NCBI Bioproject ID PRJNA1156622. Raw nanopore sequencing reads are available in the NCBI Sequence Read Archive (SRA) with Biosample accession SAMN43496284 and Run ID SRR30661341. Genomic Illumina reads are available in the SRA with Biosample accession SAMN43799130 and Run ID SRR30686038. RNAseq reads were deposited into SRA under BioProject ID SRP532563 with BioSample accession SAMN43784434 and Run ID SRR30675520. The partial COI barcode sequence for the sequenced line was uploaded to NCBI with accession number PQ361290. The *A. solani* genome is available in NCBI with accession JBHJKM000000000. The individual commands and slurm scripts as well as the KEGG assignments are available on GitHub (https://github.com/Artifice120/Aulacorthum_solani_genome_assembly_pipeline/tree/main).

## Acknowledgements

This work was supported by U.S. National Science Foundation (NSF) grant IOS-2152954 to BJP and DEB̲2305653 to PRL. BJP is a Pew Scholar in the Biomedical Sciences, supported by The Pew Charitable Trusts.

